# Defense mechanisms in *Lupinus luteus* against *Colletotrichum lupini*, involving TIR-NBS-LRR protein, hypersensitive response and phenylpropanoid pathways

**DOI:** 10.1101/2024.10.15.618496

**Authors:** Grace Armijo-Godoy, Daniela Levicoy, J. Eduardo Martínez-Hernández, Annally Rupayan, Sebastián Hernández, Makarena Carrasco, Peter DS Caligari, Riccardo Baroncelli, Haroldo Salvo-Garrido

**Author notes:** These authors contributed equally (Grace Armijo-Godoy, Daniela Levicoy).

## Abstract

Lupins are key grain legumes for future crop production, providing highly sustainable protein, essential in the face of global warming, food security challenges, and the need for sustainable agriculture. Despite their potential, lupin crops are frequently devastated by *Colletotrichum lupini*, a member of the world’s top ten fungal pathogenic genera. In our previous study, we identified *LluR1*, the first *C. lupini* resistance gene, in a wild *Lupinus luteus* accession. Further research was necessary to unravel the defense mechanisms involved. Histological analysis revealed a hypersensitive response against *C. lupini*, while transcriptome analysis highlighted a complex network of differentially expressed genes, including TIR-NBS-LRR proteins, hypersensitive response, and phenylpropanoid pathways. SNPs were identified that distinguish the protein sequences underlying immunity. These findings, along with orthology in other lupin species, offer valuable insights for developing breeding strategies to enhance *C. lupini* resistance in lupins, with significant potential impacts on food, feed, and human nutrition.

## INTRODUCTION

Grain legume crops, particularly *Lupinus* species, offer significant potential for enhancing food security and promoting sustainable agriculture (Foyer et al., 2016; Lucas et al., 2015). Traditionally used in both human and animal diets, lupins contribute to agricultural sustainability through their low fertilizer requirements and positive effects on soil fertility (Semba et al., 2021). Their versatility stems from their ability to fix atmospheric nitrogen in symbiosis with diazotrophic bacteria and solubilize phosphate in acidic soils. Among lupin species, *Lupinus luteus* stands out due to its high seed protein content and quality, making it ideal for producing plant-based proteins for food and feed (Kaczmarek et al., 2016; Schumacher et al., 2011). However, further adaptation is required for this species to meet the growing demand for plant-based proteins, particularly in light of the challenges posed by climate change, such as shifts in pathogen distribution and the emergence of new diseases. Additionally, the increased susceptibility of crops to pathogens, due to exposure to multiple stresses, represents a significant threat to food security (Abbass et al., 2022; Raza et al., 2019).

Anthracnose, caused by the fungus *Colletotrichum lupini*, is one of the most pressing threats to lupins, causing substantial yield losses, with some reports of up to 100% losses, thereby limiting production of this highly sustainable plant protein (Dubrulle et al., 2020). *C. lupini* belongs to the *Colletotrichum* genus and the *C. acutatum* species complex (Baroncelli *et al*., 2017, Damm *et al*., 2012, Talhinhas *et al*., 2016). It is a hemibiotrophic fungus that infects key lupin species such as *L. albus, L. luteus, L. angustifolius*, and *L. mutabilis* (Alkemade et al., 2021, 2022; Guilengue et al., 2022). The infection process involves three stages: following penetration, *C. lupini* initially grows subcuticularly between the cuticle and epidermal cell walls, followed by a biotrophic phase (lasting from 24 to over 72 hours), which is characterized by appressorium development and intracellular hyphae formation. Finally, the necrotrophic phase begins, during which secondary hyphae spread through host tissues (Guilengue et al., 2022; Dubrulle et al., 2020). During the early stages of infection, *C. lupini* upregulates genes involved in processes such as transcription, translation, and oxidoreduction activities, as well as encoding candidate genus-specific effectors (Dubrulle et al., 2020). As the infection progresses to the necrotrophic phase, genes encoding carbohydrate-active enzymes (CAZymes), particularly glycosyl hydrolases and pectin lyases, are upregulated. These enzymes degrade the plant cell wall, facilitating penetration and colonization. The necrotrophic phase is marked by a further increase in the expression of genes encoding pathogenicity factors, including: candidate effectors, NEPs (necrosis and ethylene-inducing proteins), and various transmembrane transporters, particularly those from the major facilitator superfamily (MFS), ATP-binding cassette (ABC) superfamily, and P-type ATPase (P-ATPase) superfamily (Guilengue et al., 2022; Dubrulle et al., 2020). These transporters are likely to play a role in nutrient acquisition and detoxification of plant defenses, further enhancing fungal proliferation.

In response to fungal pathogens, plants have evolved sophisticated defense mechanisms (Zhang et al., 2022). Without a specialized immune system, individual plant cells are responsible for recognizing and triggering defense responses (Spoel & Dong, 2012). Defense against biotrophic and hemibiotrophic pathogens, such as *C. lupini*, is primarily mediated by the hormone salicylic acid (SA), which induces reactive oxygen species (ROS) production, resulting in cell death (hypersensitive response, HR) to contain the pathogen (Cui et al., 2015). This HR also activates defense genes encoding antimicrobial proteins known as pathogenesis-related proteins (PRs) (Mishra et al., 2024). On the other hand, defense against necrotrophic pathogens is mediated by jasmonic acid (JA) and ethylene (Et), which induce the expression of protease inhibitors, defensins, chitinases, and glucanases (Li et al., 2022; Pieterse et al., 2009).

Plant immunity is orchestrated by multiple layers of receptors, including cell-surface pattern recognition receptors (PRRs) that detect pathogen-associated molecular patterns (PAMPs) or damage-associated molecular patterns (DAMPs), triggering pattern-triggered immunity (PTI) (Ngou et al., 2022). To counteract PTI, pathogens secrete effectors, some of which are recognized by intracellular receptors known as nucleotide-binding domain leucine-rich repeat receptors (NLRs) or resistance (R) proteins, leading to effector-triggered immunity (ETI) (Pieterse et al., 2009). Both PTI and ETI pathways stimulate SA biosynthesis, promoting systemic acquired resistance (SAR) (Ngou et al., 2022).

Research on anthracnose resistance in lupins, particularly *L. angustifolius*, has focused on the genetic basis of resistance, leading to the identification of the dominant resistance gene *LanR1*, which confers high resistance in the Australian cultivar Tanjil (Książkiewicz et al., 2022; Yang et al., 2004, 2012). This gene encodes an R protein involved in ETI, and transcriptomic analysis suggests that *LanR1* may enable rapid pathogen recognition, triggering HR mediated by SA and Et signaling pathways (Książkiewicz et al., 2022). However, the precise mechanisms underlying anthracnose resistance in *L. angustifolius*, particularly the role of *LanR1*, still require further investigation.

In our previous study, genetic and comparative mapping between *L. angustifolius* and *L. luteus* identified a syntenic genomic region containing the *LanR1* resistance gene. This revealed the first candidate orthologous gene for anthracnose resistance in *L. luteus* (Lichtin et al., 2020). The wild accession used in the mapping population carried the resistant allele, which we utilized to study the defense mechanisms. Through genetic, transcriptomic, and cellular analyses, we correlated the defense response with an ETI-type mechanism mediated by the R gene *LluR1*, which led to increased ROS levels and HR, preventing *C. lupini* proliferation.

## MATERIALS AND METHODS

### Plant and Fungal Material

*L. luteus* Core 98 (PI385149) (for the purposes of this work called C98), a wild accession, and Core 195 (C195, Alu*Prot*-CGNA®), a commercial cultivar from La Araucanía Region, Chile (for the purposes of this work called C195) are part of the germplasm bank of the Agriaquaculture Nutritional Genomics Center (CGNA). They were maintained under greenhouse conditions (16-h light, 23 °C and 60–75% relative humidity).

For *in vitro* tests, C98 and C195 seeds were scarified and then hydrated for 20 minutes. Next, they were washed with 70% ethanol for one minute and then with 2.5% sodium hypochlorite for one minute. Finally, they were washed three times with sterile distilled water before sowing.

*Colletotrichum lupini* was isolated from infected plants of cultivar Alu*Prot*-CGNA®, collected from different locations in the southern region of Chile. All fungi collected showed cultural and morphological features of *Colletotrichum lupini var. setosum*, as reported by Nirenberg *et al*. (2002) (Nirenberg *et al*., 2002) and therefore the genetic group as described by Dubrulle et al. (2020). They were maintained onto potato dextrose agar (PDA) at 25 °C.

### Fungal Inoculation

*C. lupini* inoculum was prepared as describe by Dubrulle et al. (2020) from a two week-old PDA culture at 25 °C. The *L. luteus* inoculations were performed with a conidial suspension of 1×10^7^ spores·mL^-1^, dropped at the base of the radicle emerging from the seed. The inoculated tissue was incubated in a moistened Petri dish at 23 ± 2 °C under 16-h photoperiod of white, fluorescent light (2000 lx). Sampling times were selected to target the first stages of the infection, the putative switch between biotrophic and necrotrophic phases and the necrotrophic phase. Uninoculated radicle emerging from the seed (prior to spore inoculation) was used as a control (T0).

### Total RNA Isolation

Approximately 150 mg of fresh hypocotyl tissue were frozen with liquid nitrogen and powdered by grinding in a mortar for transcriptomic and qRT-PCR analyses. RNA isolation was performed using Quick-RNA Plant Mini-Prep Kit (Zymo Research) following the protocol provided by the manufacturer with an extension to 30 min in DNAse digestion. RNA concentration and A260/A280 were measured by absorbance in Synergy HTX multi plate reader with Take3 Trio plate (Biotek). Integrity of RNA assessment was performed through 2% agarose gel and electrophoresis run at 100V for 1 hour.

### Transcriptome Analysis

A total of 32 RNA samples were sequenced corresponding to inoculation sampling: 0, 24, 60, 84 hours post inoculation (hpi) for both resistant and susceptible plants (n=4 biological replicates) in CD-Genomics, NY – USA in the Illumina Novaseq 6000 platform for 150 cycles in paired-end mode.

Raw RNA seq data for susceptible and resistant plants were quality control examined using FastQC v0.11.9 (https://www.bioinformatics.babraham.ac.uk/projects/fastqc/) and multiqc v1.14 (Ewels *et al*., 2016). Then, the low-quality reads were trimmed (<Q30) and Illumina adapters removed using fastp v0.23.2 (Chen *et al*., 2018). The filtered reads were mapped to the *L. luteus* reference genome (accession GCA_964019355.1) using hisat2 v2.2.1 (Kim *et al*., 2019). Gene count matrices for each experiment were calculated using feature Counts v2.0.6 (Liao *et al*., 2014). Then, the resulting count matrix was normalized and used to identify the differentially expressed genes with DESeq2 version 1.42.1(Love *et al*., 2014) All genes with an adjusted *P value* of ≤ 0.05 and absolute log2 fold change of ≥ 2 were considered differentially expressed.

For functional Gene Ontology (GO) enrichment analysis, the Cytoscape app BiNGO v3.0.5 (Pratt *et al*., 2023) in Cytoscape v3.10.1 (https://cytoscape.org/) was used.

For sequence alignment of *LluR1* orthologs, *Lupinus albus* and *Lupinus angustifolius* amino acid sequences were obtained from public databases and a multiple sequence alignment was performed using Clustal Omega (https://www.ebi.ac.uk/jdispatcher/msa/clustalo). The color-coding was based on the physicochemical properties of each amino acid was obtained using the Zappo color scheme.

### Gene Expression Analyses

The set of genes that are summarized in **Table S1** were selected and analyzed by quantitative RT-PCR. Primers were designed in Geneious Prime using Primer 3 (Koressaar & Remm, 2007) from reads obtained in the RNA-seq experiment. Inoculated samples corresponding to 0, 24, 60 and 84 hpi were analyzed, exploiting the two-step PCR protocol using iScript reverse transcription Supermix (Bio-Rad) kit for 1 μg of total RNA per sample. cDNA samples were diluted three-fold to perform qPCR. First-strand cDNA was amplified using SsoAdvanced Universal SYBR Green Supermix and CFX96 Real-Time PCR Detection System (Bio-Rad). PCR programming consisted of an initial denaturation step at 95°C for 30 seconds, 40 cycles of denaturation at 95°C for 10 seconds and annealing-extension at 60°C for 30 seconds. The specificity of reaction was tested after every PCR by melt curves (70°C-95°C). qRT-PCR reactions were run as 3 technical replications. Assessment of primer efficiency was performed using LinRegPCR software version 2021.2 (Ramakers *et al*., 2003) with efficiency values between 1.8-2.0 and R2 >0,98 considered as suitable. To find the reference gene, we tested four candidate genes: *EUKARYOTIC INITIATION FACTOR* 4a1, *EUKARYOTIC INITIATION FACTOR 1, SUCCINATE DEHYDROGENASE 1* and *PHOSPHOGLYCERATE KINASE* (Cytosolic). To select the most stable reference gene in the experiment, we used two algorithms: geNorm (an extension in Biorad CFX Maestro, (Vandesompele *et al*., 2002)) and the NormFinder R package (Andersen *et al*., 2004) selecting *EUKARYOTIC INITIATION FACTOR* 4a1 as reference gene. Calculations were performed to check the response to inoculation (i.e., expression in the inoculated samples divided by expression in control). Fold induction was calculated by normalizing relative to the housekeeping gene and control condition by the delta-delta Ct method. For statistical analysis, the normality of the data was analyzed by the Shapiro-Wilk test. If one of the samples to be analyzed showed non-parametric behavior, statistical differences were determined through the Mann-Whitney test (non-parametric statistical test equivalent to the t-test).

### Histological Assays

Hypocotyls from the susceptible and resistant plants were inoculated with *C. lupini* and harvested at 0, 24, 48, and 60 hpi, with five biological replicates. First, the samples were stained with 3,3-diaminobenzidine (DAB)-HCl to evaluate reactive oxygen species generation, according to Thordal-Christensen *et al*. (1997) (Thordal-Christensen *et al*., 1997). Then they were stained with trypan blue to evaluate fungal proliferation (Vogel & Somerville, 2000) and mounted on slides for bright-field microscopy visualization of fungal structures in a Nikon Eclipse 80i microscope (Nikon Instruments Inc., Tokyo, Japan). The plants were grown under glasshouse conditions (16 h photoperiod, 25/20°C day/night, and 70% relative humidity).

For Scanning Electron Microscopy (SEM) analysis, plant tissues inoculated with *C. lupini* were fixed in 2.5% glutaraldehyde, dehydrated, subjected to critical point drying with acetone/CO2, and then shaded with gold. This procedure was carried out by the advanced microscopy unit (UMA) of the P. Universidad Católica de Chile. The observations were made in a Hitachi TM3000 scanning microscope at 15 kV.

## RESULTS

### Differential Expression of the *LluR1* Gene to *C. lupini*

To explore *LluR1* expression during infection with *C. lupini*, resistant (C98) **(Figure 1A**) and susceptible (C195) (**Figure 1B**) plants (Lichtin et al., 2020) were inoculated. RNA isolation and qRT-PCR were performed on hypocotyls at 0, 12, 24, 60, and 84 hours post-inoculation (hpi). The results revealed a differential expression pattern between resistant and susceptible plants, with resistant plants showing significantly higher *LluR1* induction at 60 and 84 hpi (**Figure 1C**).

**Figure 1:**
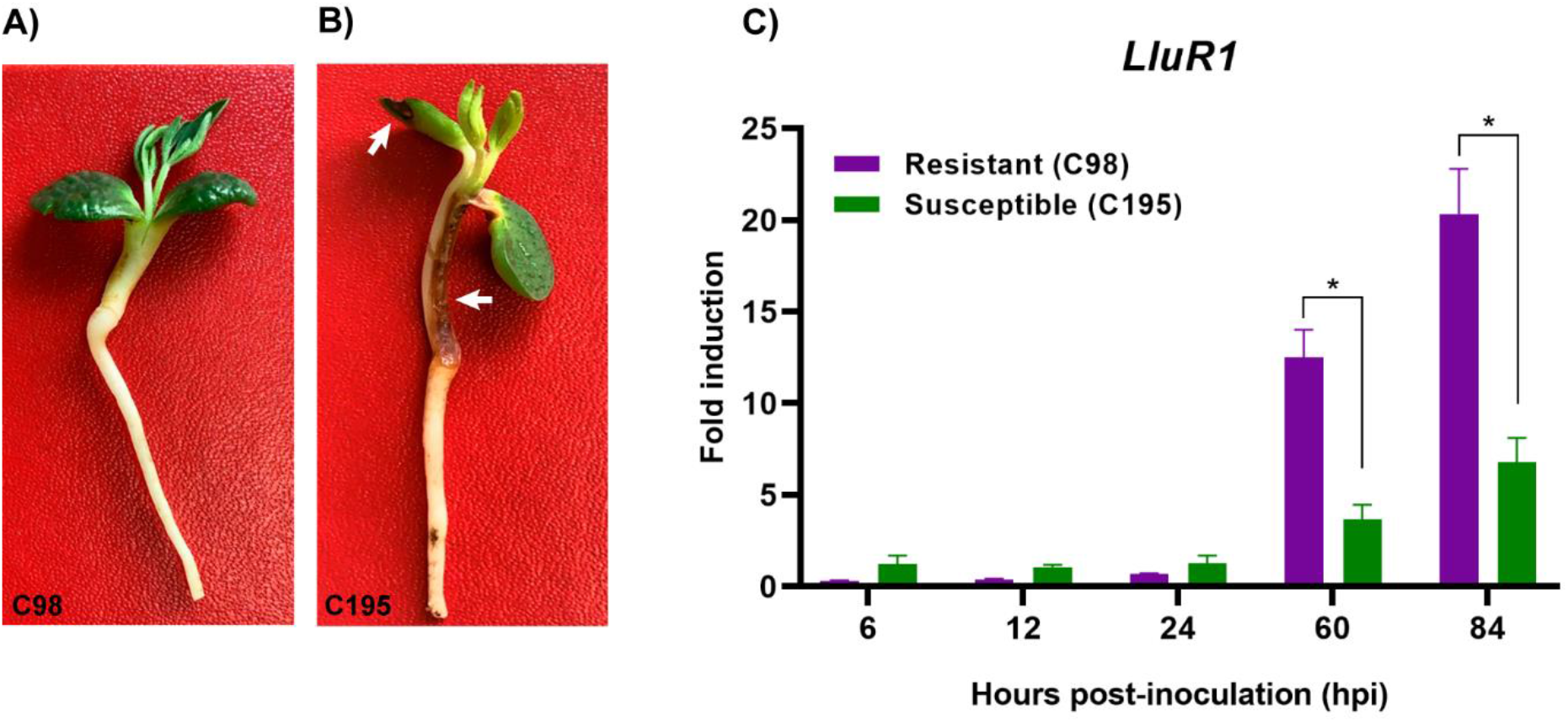
Differential Expression of the *LluR1* Gene. Resistant (A) and susceptible (B) plants, inoculated with *C lupini*. Arrows indicate disease symptoms. (C) Differential expression of *LluR1* gene in hypocotyls at 6, 12, 24, 60 and 84 hours, post inoculation with *C. lupini*. Expression levels were calculated by the delta-delta Ct method. Fold induction represent the mean ± SEM of 4 biological replicates. Statistical differences were determined through the Mann-Whitney test. * p <0, 05.

### *LluR1* encodes an R protein comprising TIR-NBS-LRR domains

The amino acid sequences of the *LluR1* gene product were studied, and functional annotation revealed that the protein encoded by *LluR1* contains a Toll/interleukin-1 receptor homology (TIR) domain at the N-terminus. This was followed by an NB-ARC and three leucine-rich repeat domains at the C-terminus **(Figure 2 and Figure S1)**, consistent with TIR-NBS-LRR resistance (R) proteins. Alignment of *Llu*R1 orthologues showed high sequence identity between Lupinus species. This demonstrates the orthology with *L angustifolius*, as proposed by us previously (Lichtin *et al*., 2020), but also newly identifies the orthology in *L albus* **(Figure 2)**.

**Figure 2:**
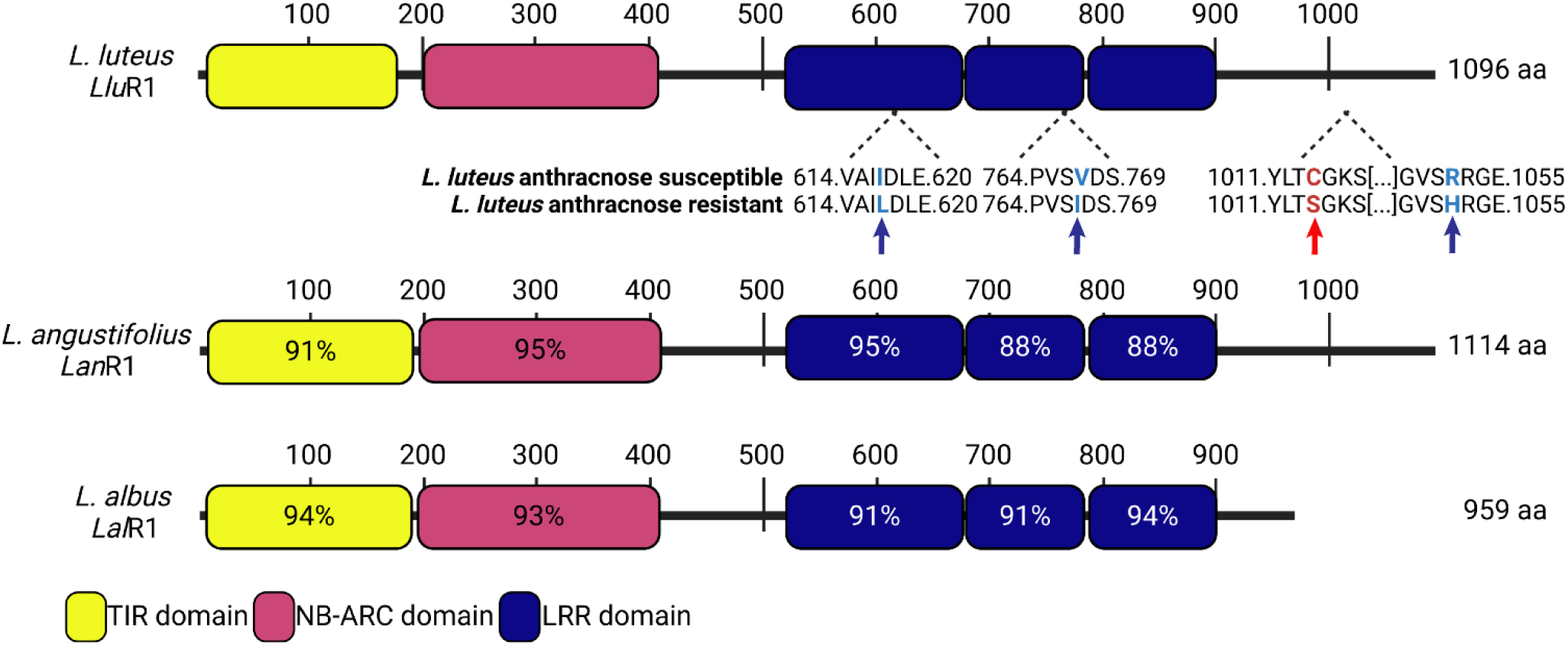
Functional annotation of *LluR1* and alignment with orthologues in *L. angustifolius* and *L. albus*. Three main domains were characterized in the amino acid sequence of *L. luteus* plant disease resistance (R) protein *LluR1* and their orthologues in related lupins: Toll/interleukin-1 receptor (TIR), NB-ARC and Leucine-rich repeat (LRR). The identity between each domain of *L. luteus LluR1* and their relatives was expressed in percentage (%). The protein sequences resulting from the SNPs in susceptible and resistant plants of *L. luteus* are displayed. The arrows identify SNPs, blue are conservative and red non-conservative (based on the physicochemical properties of each amino acid). The identity between *LluR1* and their orthologues in other lupins (*L. angustifolius* and *L. albus*) is plotted for each annotated domain.

Pairwise sequence alignments were conducted to assess single nucleotide polymorphisms (SNP) in *LluR1* gene and their effect on the amino acid sequence of resistant and susceptible plants. Four SNPs were identified, three of them resulted in conservative changes in amino acids of the coding protein: : Ile for Leu (both aliphatic/hydrophobic amino acids); Val for Ile (both aliphatic/hydrophobic amino acids), and Arg for His (both positively charged amino acids) between resistant and susceptible plants, respectively **(Figure 2)**. One SNP led to a non-conservative amino acid change (Cys, a moderately polar sulfur-containing amino acid, for Ser a polar/hydrophilic amino acid) **(Figure 2)**.

These findings suggest that *Llu*R1, *Lan*R1 and *Lal*R1 are orthologues with similar structures and functions involved in disease resistance in lupins.

### Defense response against *C. lupini* involves the activation of phenylpropanoid and salicylic acid gene pathways

To analyze the defense response against *C. lupini* at the transcriptional level, we compared the inoculated susceptible and resistant plants at 0, 24, 60, and 84 hpi. Transcriptomic analyses were conducted using 32 cDNA libraries, as described in the Methodology and **Table S2**. In the resistant plants the global expression analysis revealed a high number of differentially expressed genes (DEGs) and a gradual increase in the number of upregulated genes throughout the infection kinetics **(Figure 3A)**. While in the susceptible plants, a high number of DEGs were only observed at 24 hpi, and the upregulated DEGs decreased dramatically at 60 hpi, before rising again at 84 hpi **(Figure 3A)**.

**Figure 3:**
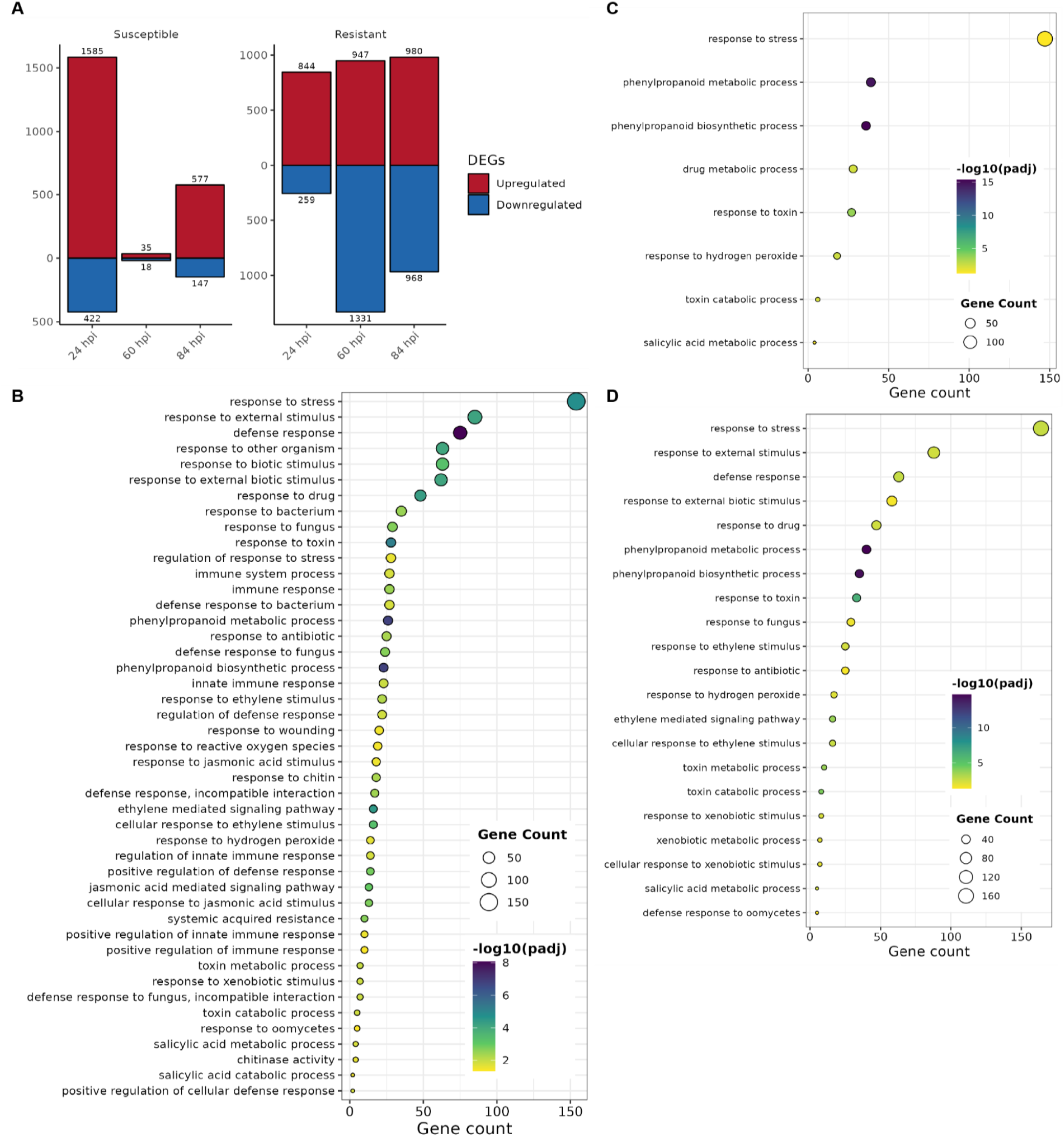
Global transcriptomic profiles and biological processes linked to response to infection. **(A)** The distribution of DEGs at the hours evaluated. The box indicates the number of DEGs downregulated (blue) and upregulated (red) at an adjusted P value of 0.05 and logFC 2. The numbers at the end of each bar correspond to the total DEGs in each plant. **(B), (C)** and **(D)** show the gene ontology and biological processes at 24, 60, and 84 hpi respectively.

Comparing the DEG categories at 24, 60, and 84 hpi in the resistant plants, significantly represented categories were found at 24 hpi including Stress response, Response to biotic stimuli, Defense, and Specific responses to ethylene, Reactive oxygen species, Jasmonic acid, Phenylpropanoids, and Salicylic acid, highlighting the breadth of defense mechanisms against *C. lupini* **(Figure 3B, Table S3)**. Interestingly, these categories declined at later hpi, while the phenylpropanoid and salicylic acid pathways remained active at 60 hpi **(Figure 3C, Table S4)** and 84 hpi **(Figure 3D, Table S5)**, while the ethylene response genes re-activated at 84 hpi **(Figure 3D, Table S5)**. Plants possess a primary or basal defense, which is induced in resistant and susceptible individuals (Ngou *et al*., 2022). To analyze this basal response, the genes expressed in the susceptible and resistant plants were compared to identify common mechanisms activated **(Figure S2)**. Interestingly, we observed common genes only at 24 **(Figure S2A)** and 84 hpi **(Figure S2C)**. Analyzing their gene ontologies, we found categories related to Response to stimuli, Stress response and Response to biotic stimuli. Response to hormones was poorly represented, and notably, the phenylpropanoid pathway was not detected **(Figure S3, Table S6)**. To analyze the secondary defense response, the genes expressed only in the resistant plants were followed. This showed that 34 common genes were upregulated at 24, 60, and 84 hpi **(Figure 4A)**. Among these, the phenylpropanoid pathway emerged as one of the most overrepresented categories, alongside more general categories such as Secondary metabolism **(Figure 4B, Table S7)**. Eight of these resistance genes were validated through qRT-PCR. 1) Chalcone synthase (*CHS)*, which encodes an enzyme involved in secondary metabolite production (Dao *et al*., 2011) **(Figure 5A)**. 2) Cytochrome P450 gene (*CYP*), coding for an enzyme involved in NADPH and O_2_-dependent hydroxylation processes, contributing to the production of secondary metabolites, antioxidants, and phytohormones (Chakraborty *et al*., 2023) **(Figure 5B)**. 3) Ferritin gene (*FT*), coding for a ubiquitous iron storage protein that regulates iron homeostasis and oxidative stress (Nguyen *et al*., 2022) **(Figure 5C)**. 4) Peroxidase gene (*PRX*), encoding an enzyme involved in various physiological processes, including defense mechanisms against pathogen infection (Kawano, 2003) **(Figure 5D)**. 5) Phenylalanine ammonia-lyase gene (*PAL*), encoding a key enzyme involved in the biosynthesis of a wide variety of specific phenylpropanoid derivatives in response to environmental factors such as pathogen infection (Barros & Dixon, 2020) **(Figure 5E)**. 6) Pathogenesis-related protein 10 gene (*PR-10*), part of a multigenic family of small acidic proteins, with RNase and DNase activity and antimicrobial and antifungal properties (Lopes *et al*., 2023) **(Figure 5F)**. 7) APETALA2/ETHYLENE RESPONSE FACTOR (*AP2/ERF*), belonging to a large superfamily induced by stress-related stimuli, such as wounding, JA, ethylene, salicylic acid, or infection by different types of pathogens (Pré *et al*., 2008) **(Figure 5G)**. 8) UDP-Glycosyltransferase (*UGT*) gene, encoding an enzyme involved in the biosynthesis of many secondary metabolites and, in many cases, regulating the activity of signaling molecules and defense compounds (von Saint Paul *et al*., 2011).

**Figure 4:**
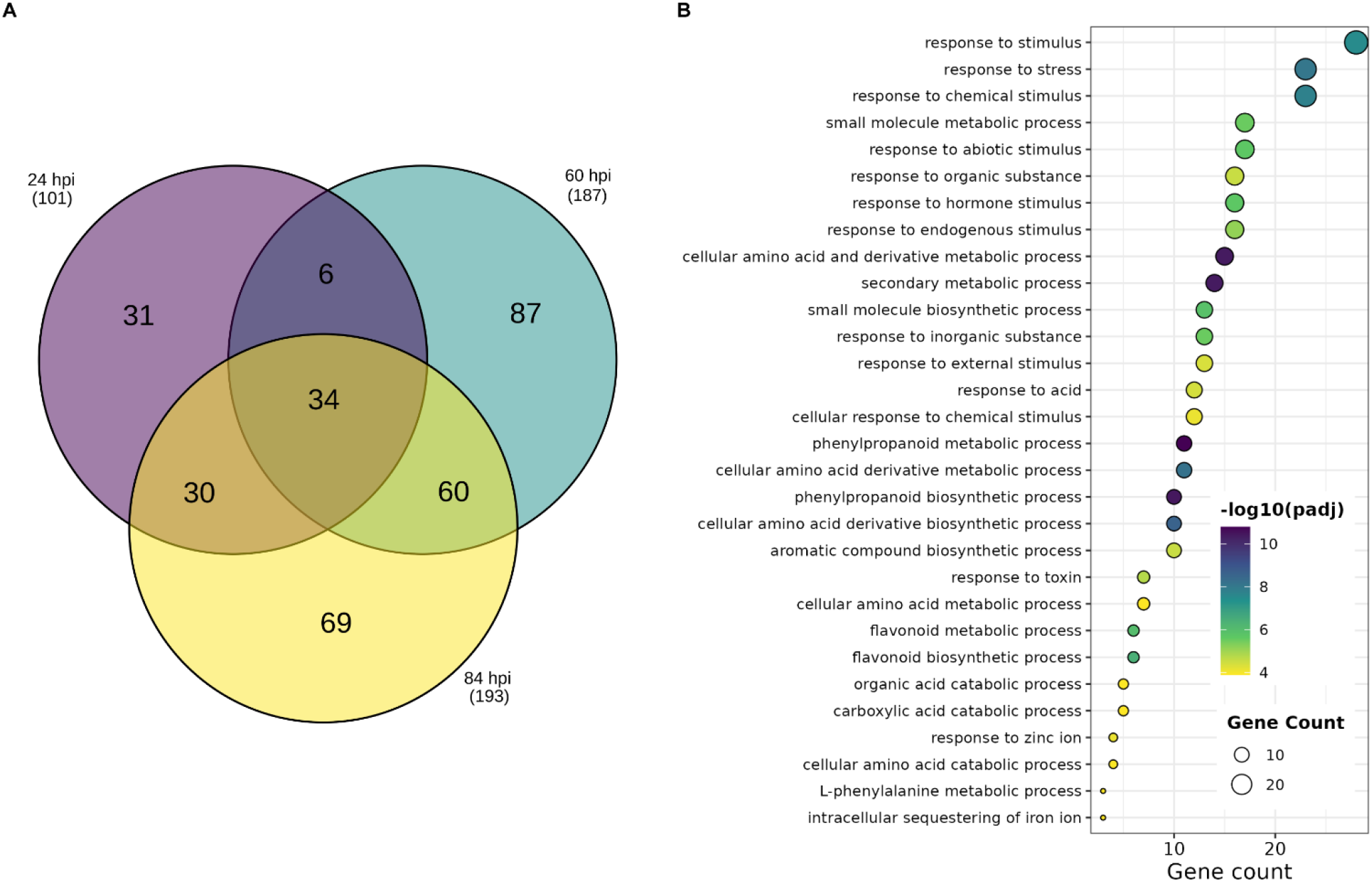
Overexpressed genes in resistant plants over different hpi. **(A)** Venn diagram illustrating differentially expressed genes (DEGs) to each hpi in resistant plants and shared across all evaluated times **(B)** GO term enrichment analysis of 34 DEGs shared across all hpi.

**Figure 5:**
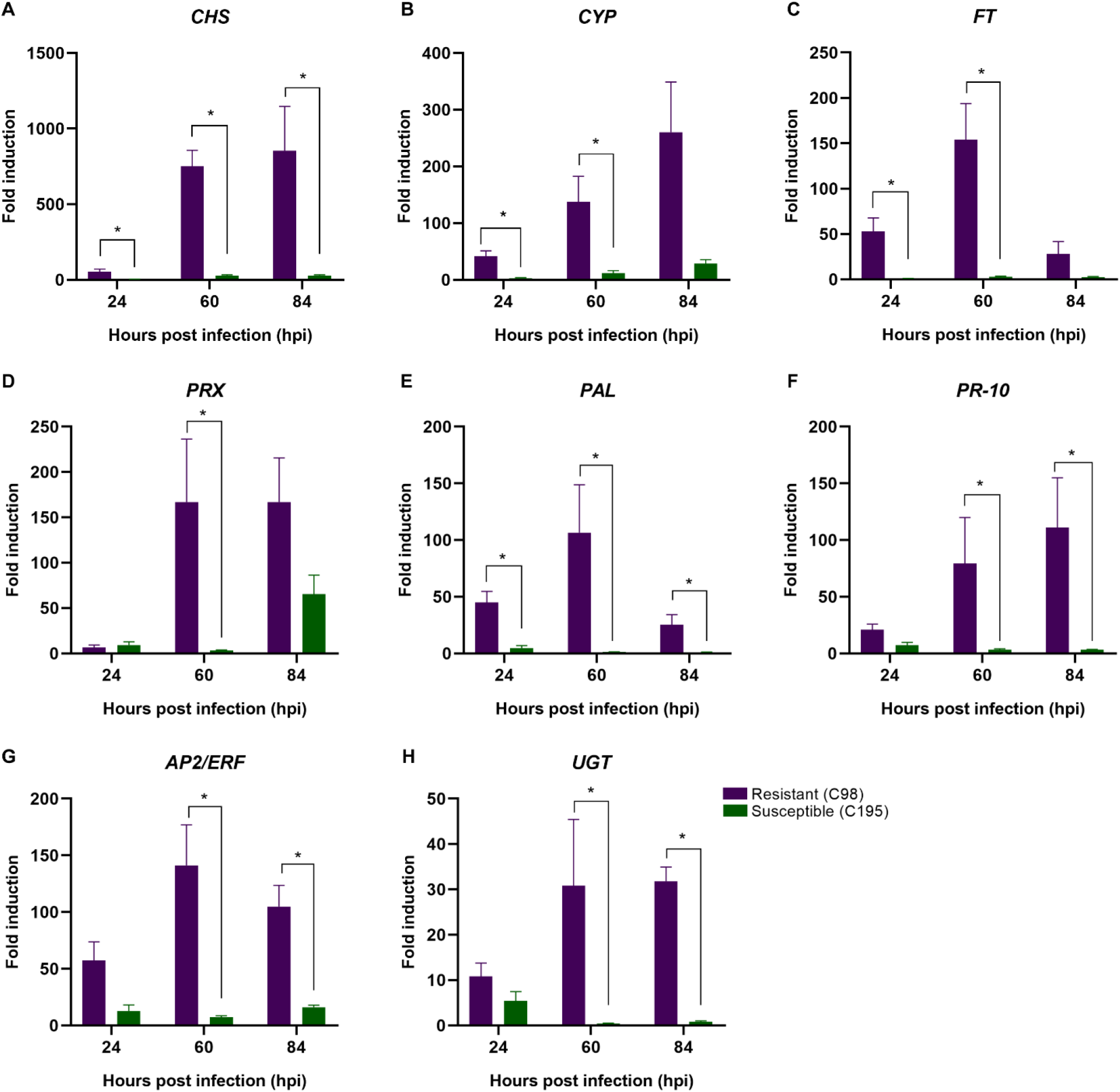
Gene expression of selected genes related to the defense mechanism. *L. luteus* plants were inoculated with *C. lupini* and sampled at 0, 24, 60, and 84 hours post inoculation (hpi). Expression levels were calculated by the delta-delta Ct method. Fold induction represent the mean ± SEM of 4 biological replicates. Resistant genotype (C98), purple bars, susceptible genotype (C195), green bars. **(A)** Chalcone synthase *(CHS)*, **(B)** Cytochrome P450 (*CYP)*, **(C)** Ferritin (*FT)*, **(D)** Peroxidase (*PRX)*, **(E)** Phenylalanine ammonia-lyase (*PAL)*, **(F)** Pathogenesis-related protein 10 (PR-10), **(G)** APETALA2/ETHYLENE RESPONSE FACTOR (AP2/ERF) and **(H)** UDP-Glycosyltransferase (UGT). For statistical analysis, the normality of the data was analyzed by the Shapiro-Wilk test, and statistical differences were determined through the Mann-Whitney test. * p <0, 05.

As expected, all these genes showed a significantly higher induction in the resistant plants, with the *CHS* gene exhibiting the highest induction rate relative to the control (T0) **(Figure 5A)**. Interestingly, *CHS* and *PAL* were activated at all evaluated times **(Figures 5A and E**), while the genes *CYP* and *FT* showed significantly higher activation at 24 and 60 hpi **(Figure 5B and 5C)**. Regarding *PRX, PR-10, AP2/ERF*, and *UGT*, they showed activation at 60 and 84 hpi **(Figures 5D, F, G, and H)**.

These results show a specific response in the resistant plants, characterized by the expression of key genes involved in secondary metabolite production and defense signaling pathways.

### The hypersensitive response generated by the resistant plants limits the fungal growth

To investigate the mechanism responsible for the defense response in the resistant plants, we conducted a histological analysis of the inoculated tissues. Hypocotyls from both susceptible and resistant plants were inoculated with *C. lupini* and then stained with 3,30-diaminobenzidine (DAB)-HCl to assess the generation of reactive oxygen species (ROS) and with trypan blue to evaluate fungal proliferation.

We examined the timing and occurrence of the oxidative burst through microscopic analysis of DAB-stained ROS generated at the site of inoculation. **(Figure 6A)**. The susceptible plants displayed rapid establishment and growth of C. *lupini* on the inoculated tissue at all evaluated times, demonstrating successful penetration, hyphae development and conidiophore formation, along with a lack of ROS containment at the infection sites. Conversely, the resistant plants exhibited ROS generation at the infection sites, restraining *C. lupini* development **(Figure 6A)**. By 24hpi, small germinating hyphae and spores were observed in the resistant plants, whereas in the susceptible plants, these hyphae were larger **(Figure 6A**, upper panel). By 48 hpi, minimal further fungal development was observed in the resistant plants, whereas in the susceptible ones, there was substantial colonization by the pathogen and uncontrolled ROS generation, associated with cell death **(Figure 6A**, middle panel). Finally, by 60 hpi, fungal development was halted in the resistant plants, with cells in contact with fungal structures exhibiting contained ROS generation associated with the hypersensitive response (HR). In contrast, in the susceptible plants, the pathogen had completed its life cycle, generating conidiophores and spores that colonized the tissue, leading to browning due to ROS generation associated with its death **(Figure 6A**, lower panel). To validate these findings, we conducted scanning electron microscopy (SEM) analysis of *C. lupini*-inoculated susceptible and resistant plants. As depicted in **Figure 6B**, the tissue surface of the resistant plants showed significantly less fungal growth compared to the susceptible plants at 24 hpi. This aligns with previous observations made via light microscopy, demonstrating that part of the resistance mechanism exhibited by the resistant plants is linked to the inhibition of pathogen’s proliferation.

**Figure 6:**
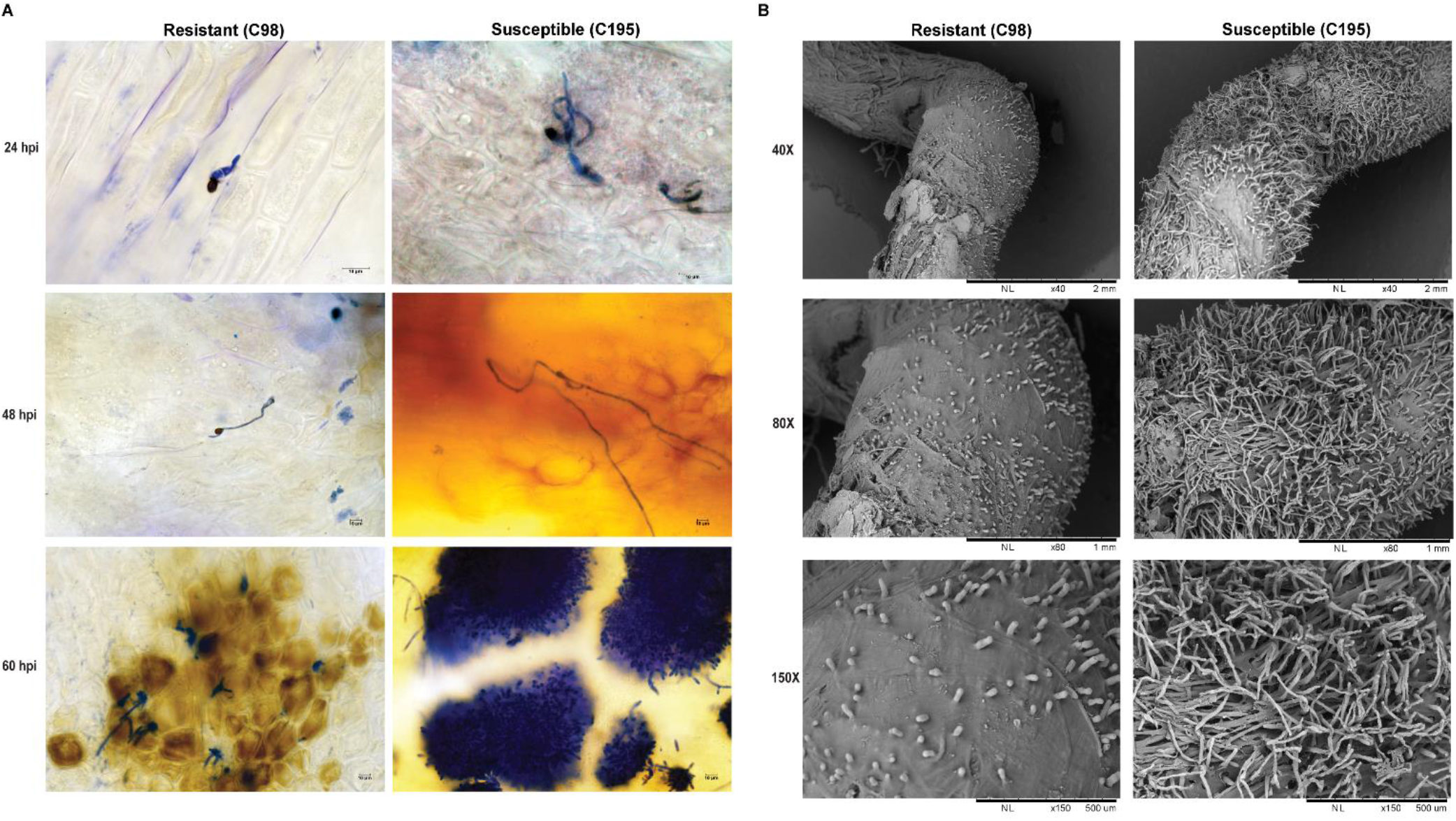
Histological analysis of susceptible and resistant plants. **(A)** *C. lupini* fungal structures (in blue) and ROS generation in *L. luteus* tissue in resistant and susceptible plants (in brown). Analysis was performed at 24, 48, and 60 hours post-inoculation. The bar represents 10 μm. **(B)** Observation of *C. lupini* fungal development in the surface of *L. luteus* tissue in resistant (C98) and susceptible (C195) plants by Scanning Electron Microscopy (SEM). Analysis was performed at 24 hpi. Magnification 40X, the bar represents 2mm; magnification 80X, the bar represents 1mm, magnification 150X, the bar represents 500 μm.

Integrating this data, it was verified that *LluR1* produces a plant R protein comprising TIR-NBS-LRR domains, with the capability to function as an immune receptor. Resistant *L. luteus*, harboring this gene, triggered a hypersensitive response and differential expression of defense genes regulated by SA and activating phenylpropanoid pathway, thus constraining the proliferation of *C. lupini*. The hypersensitive response, leading to the containment of fungal growth, thus confirmed the observations from histological analysis.

## DISCUSSION

It has been able to associate the defense response of *L. luteus* resistance to *C. lupini* with an ETI-type mechanism. The findings suggest that this mechanism is induced by the R gene *LluR1*, a *LanR1* orthologue, that codes for an R protein of TIR-NBS-LRR domains, inducing genes involved in the salicylic acid and phenylpropanoid pathways to constrain fungal proliferation. This is the first study reporting a hypersensitive response in *L. luteus* against *C. lupini*, a defense mechanism that, although conserved in plants, does not always occur, reflecting the complexity of this plant-pathogen interaction.

The dominant gene *LanR1* in *L. angustifolius* has recently been identified as responsible for high resistance to *C. lupini* in the Tanjil cultivar (Książkiewicz *et al*., 2022). In previous research, our group demonstrated the orthology of this gene with *LluR1*, marking it as the first reported gene for *C. lupini* resistance in *L. luteus* (Lichtin et al., 2020). Książkiewicz *et al*. (2022) reported an early (6 hpi) overexpression of *LanR1* (TanjilG_05042) in the 83A:476 line through RNA-seq. This pattern contrasted with subsequent qPCR analyses, where the most significant overexpression was observed in susceptible lines (Książkiewicz et al., 2022). In contrast, through RT-qPCR, we observed an upregulation of *LluR1* from 60 hpi onwards in the resistant genotype Differences in these expression patterns is currently undefined. The differential expression of the *LluR1* gene between resistant and susceptible plants inoculated with *C. lupini* reinforces the association of this gene with the resistance mechanism of *L. luteus*.

The *Llu*R1 protein contains a TIR domain in the N-terminal region, followed by a NB-ARC domain and three LRR domains in the C-terminal region, which is consistent with TIR-NBS-LRR resistance (R) proteins (Belkhadir *et al*., 2004, Lee & Yeom, 2015). These findings suggest that *Llu*R1, *Lan*R1 and *Lal*R1 are orthologues with similar structures and functions involved in disease resistance in lupins. R proteins play crucial roles in the plant immune system by directly or indirectly sensing pathogen effectors, a response known as effector-triggered immunity (ETI) (Belkhadir et al., 2004, Lee & Yeom, 2015, Pieterse *et al*., 2009). Regarding SNPs in *LluR1* of susceptible and resistant plants, even though these changes are primarily conservative, alterations in the protein sequence can impact the spatial conformation necessary for activating defense signaling (Lee & Yeom, 2015). Moreover, mutations in the R protein domains can abolish ETI, as these domains are involved in recognition, activation and downstream signaling of defense response (Elmore *et al*., 2011, Swiderski *et al*., 2009, Krasileva *et al*., 2010). Some of these mutations are located in the LRR domain, which confers recognition specificity (Ellis *et al*., 1999, Moffett *et al*., 2002) and may alter the effectiveness of recognition in the susceptible plant.

Transcriptomic analysis of inoculated susceptible and resistant plants of *L. luteus* revealed dynamic changes in gene expression patterns during infection. The resistant plants exhibited a robust defense response characterized by the activation of phenylpropanoid and salicylic acid pathways. Interestingly, the phenylpropanoids pathway is involved in the biosynthesis of many antimicrobial compounds and is one of the biosynthetic routes of salicylic acid in response to pathogen infection (Lefevere *et al*., 2020). This result is partly consistent with the transcriptomic analysis carried out by Książkiewicz *et al*. (2022) in *L. angustifolius*, where an increase in SA and ethylene signaling pathways were detected (Książkiewicz et al., 2022).

Identifying candidate resistance genes differentially expressed exclusively in the resistant plants and, subsequent qRT-PCR validation showed a specific response against *C. lupini*. Among these genes, chalcone synthase (CHS), phenylalanine ammonia-lyase (PAL), UDP-glycosyltransferase (UGT), and cytochrome P450 (CYP) were involved in the biosynthesis of flavonoids, as part of the phenylpropanoid biosynthetic pathway (Dong & Lin, 2021, Yonekura-Sakakibara *et al*., 2019). These secondary metabolites play roles in many plant functions, including defense against pathogens (Aboody & Mickymaray, 2020). They can act as signal molecules and ROS scavengers, and they also exhibit antifungal mechanisms, such as disrupting the plasma membrane, inducing mitochondrial dysfunctions, and inhibiting cell wall formation, cell division, and RNA and protein synthesis (Aboody & Mickymaray, 2020, Dias *et al*., 2021). It has been previously reported that the accumulation of flavonoids is essential for increased tolerance to biotic stress. For example, rice plants with increased accumulation of flavonoids showed greater tolerance to bacterial leaf blight disease (Jan *et al*., 2021, Lu *et al*., 2017), and flavonoids also played a role in powdery mildew resistance in wheat (Xu *et al*., 2023).

PAL and peroxidases (PRX) can participate in the fortification of the cell wall through lignification and promoting cell wall strengthening, thus making pathogen invasion more difficult (Ninkuu *et al*., 2023, Ninkuu *et al*., 2022, Wakabayashi *et al*., 2012). It has been reported that PAL induces the accumulation of lignin, SA, cinnamic acid, and fatty acid in defense against pathogens (Ninkuu et al., 2023). PRXs produce phenolic compounds and fortify cell walls through lignin formation. They also generate ROS and can participate in wound healing, playing an important role in plant defense (Kawano, 2003, Ninkuu et al., 2023).

Within the differentially expressed genes showing a higher induction in the resistant plants, ferritin (FT) stores iron and regulates its homeostasis (Zielińska-Dawidziak, 2015) and iron balance is crucial for defense against pathogen-induced oxidative stress. During pathogen infection, there is a competition between the host and the pathogen for iron (Sánchez-Sanuy *et al*., 2022). In addition, the host plant might use the toxicity of iron to reduce the proliferation of the invading pathogen (Herlihy *et al*., 2020, Sánchez-Sanuy et al., 2022). Histochemical analysis of *Magnaporthe oryzae*-infected rice leaves revealed colocalization of iron and reactive oxygen species in cells located near fungal penetration sites in plants exposed to iron (Dangol *et al*., 2019). Moreover, transgenic tobacco plants expressing alfalfa ferritin exhibited tolerance to necrotic damage caused by *Alternaria alternata* and *Botrytis cinerea* infections (Deák *et al*., 1999).

In addition to the above mechanisms, transcription factors such as AP2/ERF play a pivotal role in regulating the expression of genes involved in both abiotic and biotic stress responses, including pathogen defense (Ma *et al*., 2024, Xie *et al*., 2022). ERF gene family are involved in ethylene signaling and defense gene regulation (Xie et al., 2022). Ethylene is an essential phytohormone for plant growth, development, and stress tolerance, and the AP2/ERF transcription factors regulate its biosynthesis (Xie et al., 2022, Ma et al., 2024). AP2/ERFs act downstream of mitogen-activated protein kinase (MAPK) cascades, regulating the expression of genes associated with hormonal signaling pathways, biosynthesis of secondary metabolites, and formation of physical barriers involved in defense responses (Ma et al., 2024). Interestingly, it has been reported that MAPK in rice, phosphorylates and enhances the binding affinity of an AP2/ERF transcription factor to the promoter of PR genes, resulting in enhanced disease resistance (Cheong *et al*., 2003). Additionally, AP2/ERFs have also been reported to regulate plant resistance through the SA biosynthesis or signaling pathway (Ma et al., 2024, Wang *et al*., 2020, Zheng *et al*., 2019), JA/ET signaling pathway (Huang *et al*., 2022, Ma et al., 2024), or mediating crosstalk between SA and JA/ET signaling pathways (Ma et al., 2024).

The pathogenesis-related protein 10 (PR-10) gene was also differentially expressed exclusively in the resistant genotype. PR-10 proteins are a family related to plant response to pathogens. They have ribonuclease activity and are induced in response to infection, participating in pathogen RNA degradation and defense signaling (Agarwal & Agarwal, 2014). Interestingly, PR-10 proteins can direct carbon flux through the flavonoid biosynthetic pathway (Dastmalchi, 2021). They interact with biological ligands, including phytohormones, proteins, fatty acids, amino acids, phenolics, and several classes of alkaloids (Morris *et al*., 2021, Dastmalchi, 2021), and it has been reported in some cases that its expression is correlated with flavonoid accumulation and phenylpropanoid pathway genes (Dastmalchi, 2021, Strömvik *et al*., 1999, Warner *et al*., 1994).

Altogether, the activation of these genes suggests a robust defense response in the resistant plants of *L. luteus* against *C. lupini*, including the biosynthesis of antimicrobial compounds via the phenylpropanoid pathway, production of SA and PR proteins, transcriptional regulation by AP2/ERF, iron management, and ROS generation. These responses strengthen physical barriers, inhibit pathogen growth, and coordinate defensive signaling to protect the plant. It has been reported that *L. angustifolius* plants infected with *C. lupini* accumulate secondary metabolites like isoflavone aglycones and glycoconjugate (Wojakowska *et al*., 2013). Furthermore, *L luteus* inoculated with *Fusarium oxysporum* stimulates the phenylpropanoid metabolism, increasing the isoflavonoids concentration, as a part of the defense system of legumes (Morkunas *et al*., 2005).

The *C. lupini* -resistant *L. luteus* plants demonstrated an upregulation of genes associated with the oxidative burst, a rapid production of ROS that serves as a key indicator of plant defense mechanisms. The precise coordination of ROS production with antioxidant systems is vital, as it prevents cellular damage while ensuring effective pathogen defense (Dumanović *et al*., 2021), which is related to a hypersensitive response. Our histological analysis supports this hypothesis. Resistant plants showed ROS generation at the infection sites only in the cells in direct contact with the pathogen and the limitation of fungal development. At the same time, susceptible plants exhibited a faster establishment and development of the fungus over tissue surface in all the infection times, showing successful penetration and hypha development with conidiophore formation and the absence of ROS containment at the infection sites. Scanning electron microscopy studies further supported these findings, showing reduced fungal growth on the tissue surface of resistant plants compared to susceptible ones.

The temporality and occurrence of the oxidative burst generated at the infection site are critical for pathogen containment (Wojtaszek, 1997) and the SA-induced hypersensitive response has been shown to induce the generation of ROS early in the infection. However, this hormone stimulates the later expression of genes coding for antioxidant proteins to contain the oxidative burst and trigger cell death only in the infected tissue (Herrera-Vásquez *et al*., 2015). Initial levels of ROS would also function as signaling molecules to neighboring cells (Waszczak *et al*., 2018).

Based in this study, it is clear that the presence of *Llu*R1 as an R protein can “sense” a fungal effector, either directly or indirectly (Dangl & Jones, 2001, Van der Biezen & Jones, 1998). In plants generally there are two main pathways of defense against pathogens, where defense against biotrophic/hemibiotrophic is attributed to SA and against necrotrophic to JA (Pieterse et al., 2009). *C. lupini* has been described as a hemibiotrophic pathogen (Guilengue *et al*., 2022). In the first stage of infection, the fungus behaves as a biotrophic and develops an appressorium to penetrate the host cuticle and epidermal cells. However, appressorium formation can occur during infection due to spores that may germinate later (Dubrulle *et al*., 2020). The recognition of a *C. lupini* effector by the R-protein *Llu*R1 would activate the signaling pathway mediated by the SA hormone in this first stage of infection. Regarding the *C. lupini* infection, at 48 and 60 hpi, the observation of secondary hyphae indicates the beginning of host penetration in the hypocotyls infection (Dubrulle et al., 2020). This would be consistent with the induction of the *LluR1* gene at 60 and 84 hpi, where the pathogenic effector introduced to the tissue would be recognized, including those spores that might take longer to complete this invasion phase.

This work supports a defense response mechanism where a fungal effector is recognized by the R protein *Llu*R1, activating the ETI response. This mechanism intricately integrates components and metabolic pathways that are not unique to *L. luteus* but are shared broadly across plant species. Transcriptional regulation by AP2/ERF, activated by MAPK cascades, plays a pivotal role in the proposed defense mechanism. It regulates the biosynthesis of secondary metabolites, effectively coordinating defense responses through the SA and JA/ET pathways, which are central to plant immunity across diverse taxa. The activation of these defense pathways leads to the activation of the phenylpropanoid pathway and the biosynthesis of antimicrobial compounds, such as flavonoids. PR-10 inhibits pathogen growth and modulates flavonoid biosynthesis, integrating chemical defense and signaling. Additionally, the production of phenolic acids and lignin serves to fortify the cell wall, adding another layer of defense. Ferritin contributes to ROS formation at infection sites, helping limit pathogen spread. Finally, peroxidases are involved in lignification and ROS generation. The increase of ROS in the infected cell triggers the hypersensitive response, which generates programmed cell death at the site of infection, containing and preventing pathogen progression **(Figure 7)**. This cascade of events underlines what is basically a highly conserved defense strategy among plants, reflecting the evolutionary pressure to develop robust mechanisms to counteract pathogenic attack.

**Figure 7:**
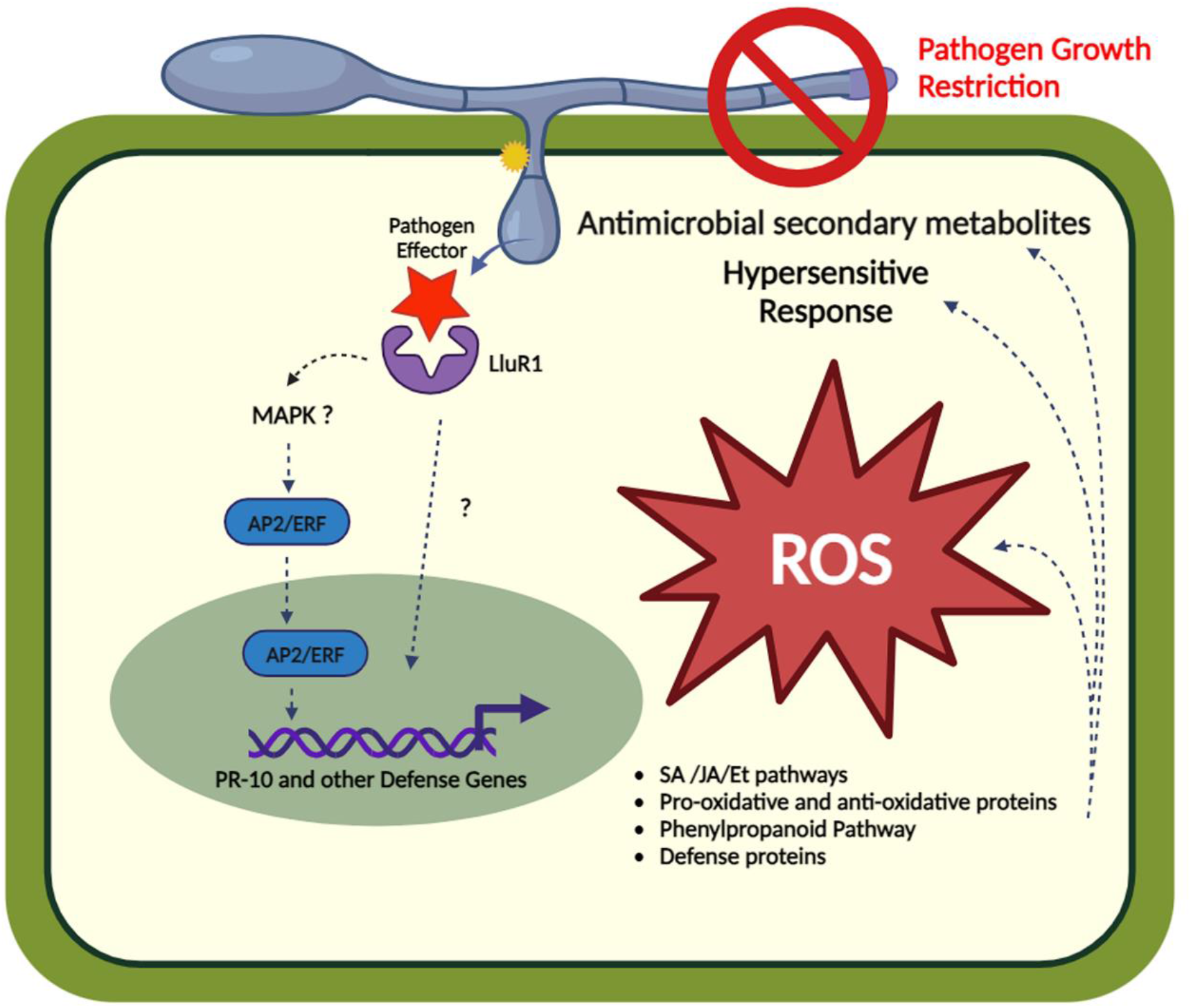
Model of the defense mechanism. A fungal effector is recognized by the R protein of LluR1, which activates various metabolic pathways. AP2/ER, activated by MAPK cascades, regulates the biosynthesis of secondary metabolites, coordinating defense responses through phenylpropanoid and SA, JA/Et pathways, leading to ROS generation and a general hypersensitive response, containing the infection and fungal development.

In conclusion, we successfully elucidated the defense response mechanism of *L. luteus, which* exhibits high resistance to *C. lupini, by* an effector-triggered immunity (ETI) mechanism. Our findings indicate that this mechanism is induced via the R gene *LluR1*, which encodes an R protein with TIR-NBS-LRR domains. The upregulation of *LluR1* in resistant plants highlights its role in activating defense responses against *C. lupini* related to the phenylpropanoid and salicylic acid pathways. These findings underline the complex and multi-faceted nature of the plant defense response in *L. luteus*, providing insights into the molecular mechanisms of resistance against *C. lupini*. This study provides valuable insights into the molecular mechanisms supporting disease resistance in *L. luteus*, with significant implications for breeding programs to enhance crop resilience.

## Acknowledgments

This research was supported by the Agri-aquaculture Nutritional Genomic Center (CGNA), ANID Centros de Investigacion Asociativa, Project R20F0003, Chile.

## Author contributions

G.A.G, D.L., A.R., S.H., M.C. and H.S.G. performed experiments. J.E.M performed bioinformatics analysis. G.A.G, D.L., J.E.M and H.S.G. designed experiments. H.S.G supervised the study and analyzed the results. P.D.S.C. and R.B. collaborated in experimental design and troubleshooting. G.A.G, P.D.S.C, and H.S.G. wrote the manuscript.

## Data availability

Raw RNA-seq reads data have been deposited at SRA NCBI as Bioproject http://www.ncbi.nlm.nih.gov/bioproject/1169352 and are publicly available as of the date of publication. Any additional information required to reanalyze the data reported in this paper is available from the lead contact upon request.

