## Supplemental Figures for "Defense mechanisms in *Lupinus luteus* against *Colletotrichum lupini*, involving TIR-NBS-LRR protein, hypersensitive response and phenylpropanoid pathways"

### Supplementary Figures

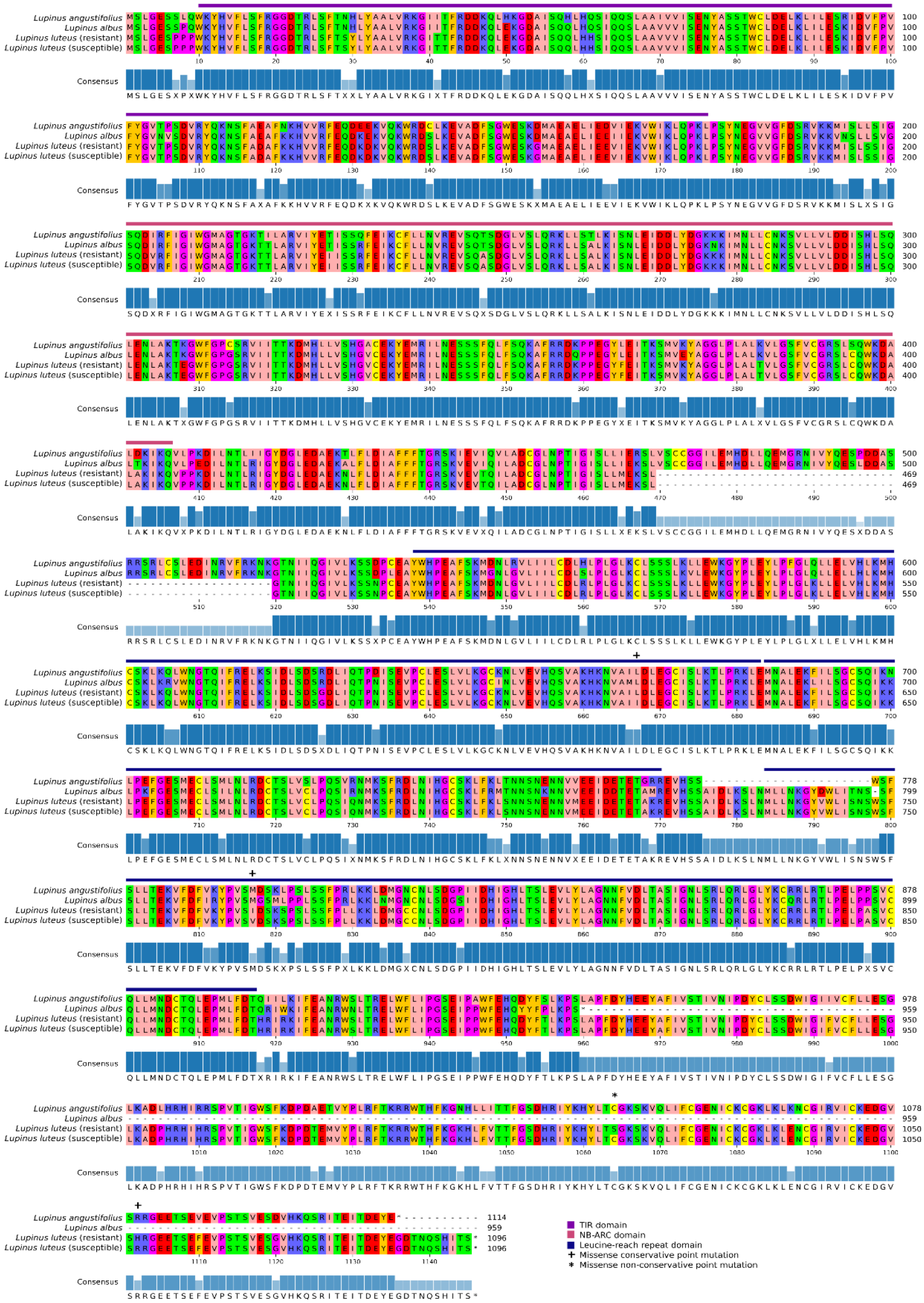

**Figure S1:** Sequence alignment of LluR1 orthologues in three lupin species, *Lupinus angustifolius*, *Lupinus albus*, and *Lupinus luteus*. The alignment is color-coded based on the physicochemical properties of each amino acid. Point mutations identified between *L. luteus* susceptible and resistant to *C. lupini* infection are indicated with a cross or asterisk, depending on whether it is a conservative mutation or non-conservative, respectively. Predicted functional domains are delineated: the TIR domain is highlighted in magenta, the NB-ARC domain in fuchsia, and the Leucine-rich repeat domain in blue.

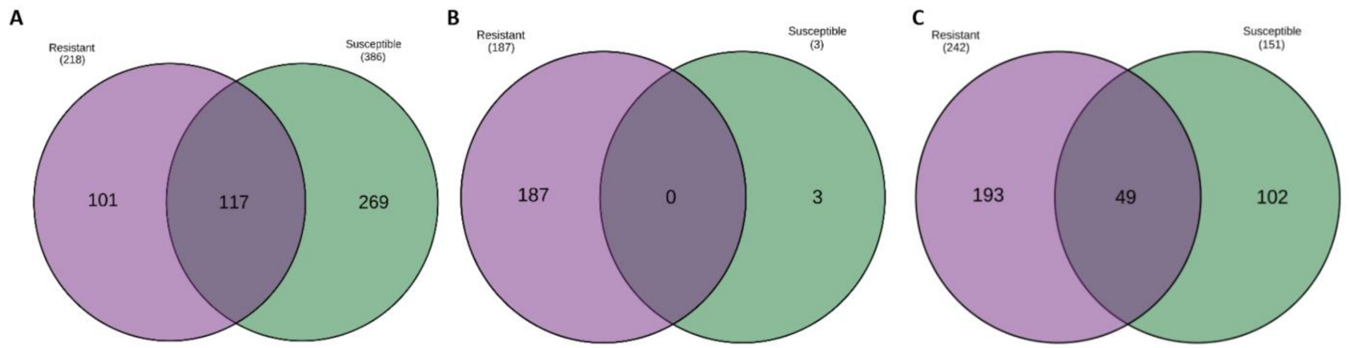


**Figure S2:** Venn diagram depicting the overlap of differentially expressed genes between resistant (purple) and susceptible (green) plants at post-inoculation times. **(A)** 24hpi. **(B)** 60 hpi. **(C)** 84 hpi.

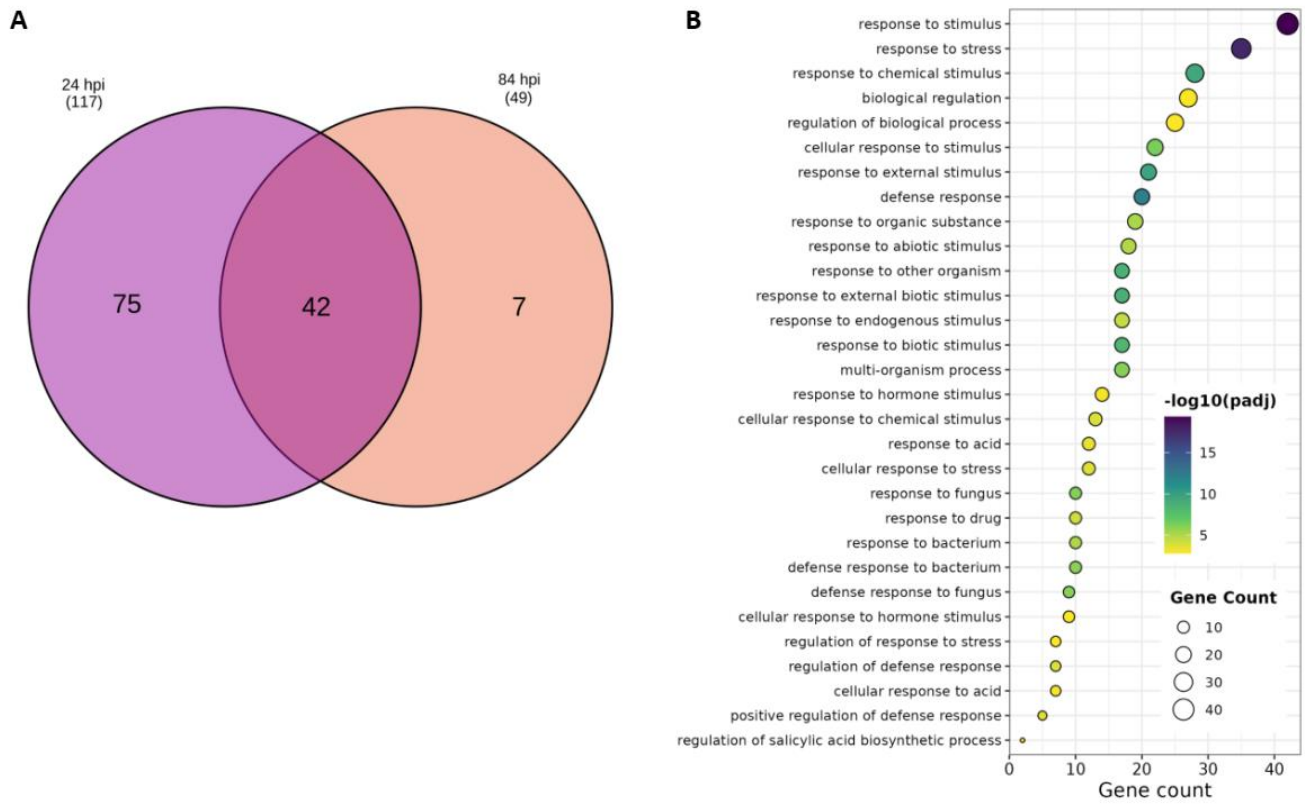


**Figure S3:** Genes associated with the immune response in *L. luteus* display distinct expression patterns in susceptible and resistant plants inoculated with *C. lupini*. **(A)** A Venn diagram illustrates the overlap of genes linked to immune response and defense in resistant and susceptible plants. **(B)** Gene Ontology (GO) term enrichment analysis highlights biological processes related to the response to *C. lupini* infection in resistant and susceptible plants.
